# Phosphorylation of phase-separated p62 bodies by ULK1 activates a redox-independent stress response

**DOI:** 10.1101/2022.12.21.521356

**Authors:** Ryo Ikeda, Daisuke Noshiro, Hideaki Morishita, Shuhei Takada, Shun Kageyama, Yuko Fujioka, Tomoko Funakoshi, Satoko Komatsu-Hirota, Ritsuko Arai, Elena Ryzhii, Manabu Abe, Tomoaki Koga, Mitsuyoshi Nakao, Kenji Sakimura, Arata Horii, Satoshi Waguri, Yoshinobu Ichimura, Nobuo N Noda, Masaaki Komatsu

## Abstract

NRF2 is a transcription factor responsible for antioxidant stress responses that is usually regulated in a redox-dependent manner. p62 bodies formed by liquid-liquid phase separation contain Ser349-phosphorylated p62, which participates in the redox-independent activation of NRF2. However, the regulatory mechanism and physiological significance of phosphorylation remain unclear. Herein, we identify ULK1 as a kinase responsible for phosphorylation of p62. ULK1 co-localizes with p62 bodies, and directly interacts with p62. This phosphorylation allows KEAP1 to be retained within p62 bodies, activating NRF2. *p62*^*S351E/+*^ mice are phosphomimetic knock-in mice in which Ser351 corresponding to human Ser349 is replaced by Glu. These mice, but not phosphodefective *p62*^*S351A/S351A*^ mice, exhibit NRF2 hyperactivation and growth retardation, the latter caused by malnutrition and dehydration due to obstruction of the esophagus and forestomach secondary to hyperkeratosis. *p62*^*S351E/+*^ mice are a phenocopy of systemic *Keap1*-knockout mice. Our results expand our understanding of the physiological importance of the redox-independent NRF2 activation pathway and provide new insight into the role of phase separation in this process.

## Introduction

Liquid-liquid phase-separated biomolecular condensates, liquid droplets play an important role in many biological processes, such as gene expression, protein translation, stress response, and protein degradation, by incorporating a variety of RNA and client proteins into their interior depending on the intracellular context ^1^. Autophagy is involved in the degradation of several cytoplasmic liquid droplets, including stress granules and P bodies, and defects in this process are thought to cause transition of these droplets to the solid phase, resulting in the development of intractable diseases such as neurodegenerative disorders and cancer ^2,3^. Of the droplets that have a unique biological function and are degraded by autophagy, p62 bodies (also called p62 droplets) are liquid droplets formed by liquid-liquid phase separation (LLPS) of p62 and its binding partners, ubiquitinated proteins ^4,5^. p62 bodies are involved in the regulation of intracellular proteostasis through their own autophagic degradation, and also contribute to the regulation of the major stress-response mechanism by sequestration of a client protein, kelch-like ECH-associated protein 1 (KEAP1) ^6,7^.

An Unc-51-like kinase 1 (ULK1) phosphorylates p62 at Ser407, inhibiting dimer formation of the UBA domain of p62 ^8,9^, and subsequent phosphorylation of Ser403 by TBK1, CK2, TAK1, and ULK1 allows binding of ubiquitinated proteins ^8,10-12^. These phosphorylation events are thought to promote LLPS ^4,5^. In the degradation of p62 bodies, the ULK1 protein kinase complex consisting of FIP200/RB1-inducible coiled-coil protein 1 (hereafter as FIP200), ULK1, ATG13, and ATG101 is translocated onto the bodies by binding of the FIP200 Claw domain to p62 ^13^. Alternatively, the ULK1 protein kinase complex is recruited to p62 bodies through the interaction of FIP200 with TAX1BP1, which localizes at p62 bodies through the interaction with a p62 binding partner NBR1 ^14^. Subsequently, ATG proteins assemble around the bodies ^15^. In the end, the p62 bodies are surrounded by autophagosomes due to the wetting effect ^16^ and the binding of LC3 or GABARAP to p62 on the isolation membrane ^15^, followed by lysosomal degradation.

KEAP1 is an adaptor protein of cullin 3 ubiquitin ligase for nuclear factor (erythroid-derived 2)-like 2 (NRF2), which is a key transcription factor for a series of genes encoding anti-oxidative proteins and enzymes ^17^. In the canonical pathway, KEAP1 is inactivated by the modification of oxidants, and NRF2 is then activated via redox-dependent regulation ^17^. This redox-dependent pathway has been shown to be important in redox, metabolism, and protein homeostasis, as well as in the regulation of inflammation and cellular protection against many pathological conditions ^17,18^. In addition to this canonical pathway, a specific region of p62 directly interacts with KEAP1, competitively preventing the interaction between KEAP1 and NRF2 ^19^. The phosphorylation of Ser349 locating at the KEAP1-interacting region of p62 enhances the interaction of p62 with KEAP1, sufficiently resulting in full activation of NRF2 independently of redox conditions ^20^. However, the kinase(s) and regulatory mechanism underlying the redox-independent pathway, as well as its physiological significance, remain unclear.

Herein, we show for the first time that ULK1 is a major kinase for Ser349 of p62, both *in vitro* and *in vivo*. ULK1 directly interacts with p62 and phosphorylates Ser349 of p62. ULK1 localizes in p62 bodies *in vitro* and *in vivo* in a FIP200-independent fashion. While this phosphorylation does not affect the influx of KEAP1 into p62 bodies, it inhibits KEAP1 outflow, keeping KEAP1 in the p62 bodies and activating NRF2. Knock-in mice with a phosphomimetic mutation, but not those with a phosphodefective mutation, exhibit persistent activation of NRF2, which causes hyperkeratosis and consequently obstruction of the esophagus and forestomach, and eventually severe growth retardation due to malnutrition. Taken together, these results indicate the physiological importance of p62 body- and ULK1-dependent and redox-independent stress response.

## Results

### ULK1 directly interacts with and phosphorylates p62

To clarify whether the UKL1 kinase itself has an effect on physical property and physiological role of p62 bodies, we first studied physical interaction of p62 with ULK1 or its yeast homologue Atg1 using high-speed atomic force microscopy (HS-AFM) (Fig. 1). ULK1 has a serine-threonine kinase domain (KD) at the N-terminus and two microtubule interaction and transport (MIT1 and 2) domains at the C-terminus, all of which are conserved between yeast and mammals (Fig. 1a). The KD and MIT1/2 domains are linked by an intrinsically disordered region (IDR) (Fig. 1b). p62 has an N-terminal Phox1 and Bem1p (PB1) domain and C-terminal ubiquitin-associated (UBA) domain as well as several interacting regions such as LIR and KIR located in an IDR between the PB1 and UBA domains (Fig. 1a and b). We purified recombinant p62 (268–440 aa and 320–440 aa) and SNAP-tagged ULK1 and Atg1 (Fig. 1c). HS-AFM of SNAP-ULK1 revealed that it contains two globular domains consisting of a KD and two tandem MIT domains, which are linked to each other with an IDR, like Atg1 ^21^ (Supplementary Fig. S1a and Supplementary Movie S1). Meanwhile, HS-AFM of p62 (268–440 aa) visualized the homodimeric structure mediated by the dimerization of the UBA domain, forming a hammer-shaped structure with IDRs wrapped around each other (Supplementary Fig. S1b and Supplementary Movie S2). When each SNAP-ULK1 and SNAP-Atg1 was mixed with p62 (268–440 aa), the p62 homodimer directly bound to SNAP-ULK1 and SNAP-Atg1 via dynamic IDR-IDR and IDR-globular domain interactions (Fig. 1d and e, Supplementary Fig. S1c, and Supplementary Movie S3 and 4). Consistent with this, ULK1 and Atg1 directly phosphorylated recombinant p62 (268– 440 aa and 320–440 aa) at Ser349 (Fig. 1f). Although Ser403 was hardly phosphorylated (Fig. 1f), it was also phosphorylated when mCherry-tagged full-length p62 was used (Fig. 1f), indicating that the N-terminal PB1 domain of p62 is required for efficient Ser403 phosphorylation by ULK1 and Atg1. These data suggest that ULK1 directly interacts with and phosphorylates p62.

**Fig. 1.**
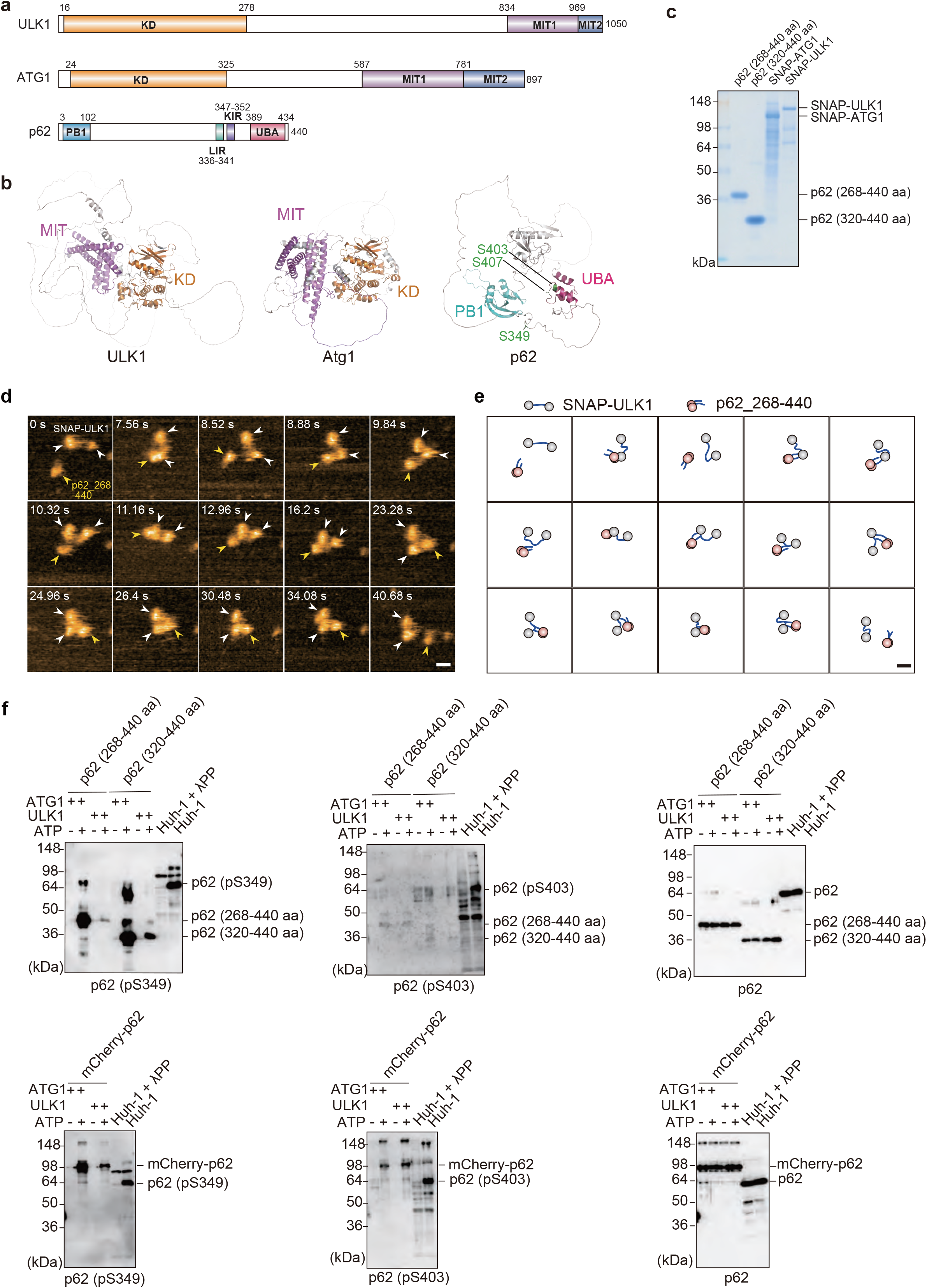
Molecular dynamics of ULK1 and p62. **a** Domain structures of ULK1, Atg1, and p62. KD, kinase domain; MIT, microtubule interaction and transport domain; PB1, Phox1 and Bem1p domain; LIR, LC3-interacting region; KIR, KEAP1-interacting region; UBA, ubiquitin-associated domain. **b** Predicted three-dimensional structures of ULK1, Atg1, and p62 by Alphafold 2. **c** CBB staining of purified p62 (268–440 aa), p62 (320–440 aa), SNAP-Atg1, and SNAP-ULK1. **d** Successive HS-AFM images of p62_268–440 with SNAP-ULK1. Height scale: 0–3.5 nm; scale bar: 20 nm. **e** Schematics showing the observed molecular characteristics by HS-AFM. Grey spheres, globular domains consisting of N-terminal KD and C-terminal MIT domain of ULK1; pink spheres, globular domains consisting of C-terminal UBA domain of p62; blue thick solid lines, IDRs. **f** *In vitro* kinase assay. Purified recombinant p62 (268–440 aa), p62 (320–440 aa) or mCherry-p62 was incubated for 20 min at 30 °C with purified SNAP-Atg1 or SNAP-ULK1 in the presence or absence of ATP. Reactions were then terminated by adding LDS sample buffer containing reducing agent, followed by immunoblot analysis with the indicated antibodies. As positive and negative controls, Huh-1 cell lysates treated with or without lambda protein phosphatase (λPP) were used. Data were obtained from three independent experiments.

### Localization of ULK1 in p62 bodies

p62 undergoes LLPS upon interaction with ubiquitinated proteins *in vitro*, forming p62 condensates ^4^. We examined whether SNAP-Atg1 and SNAP-ULK1 associate with p62 condensates *in vitro*. Consistent with previous reports ^4,5,15^, mixing mCherry-p62 with linear octa-ubiquitin (8xUb) resulted in the formation of condensates (Fig. 2a). Their size was increased by phosphomimetic p62 mutations (S403E and S407E) (Fig. 2a) that are known to increase the binding affinity of p62 to ubiquitin ^8,10,11^. SNAP-Atg1 and SNAP-ULK1 were recruited to p62 condensates when all were incubated together (Fig. 2b), implying that both Atg1 and ULK1 associate with p62 even in the droplet form.

**Fig. 2.**
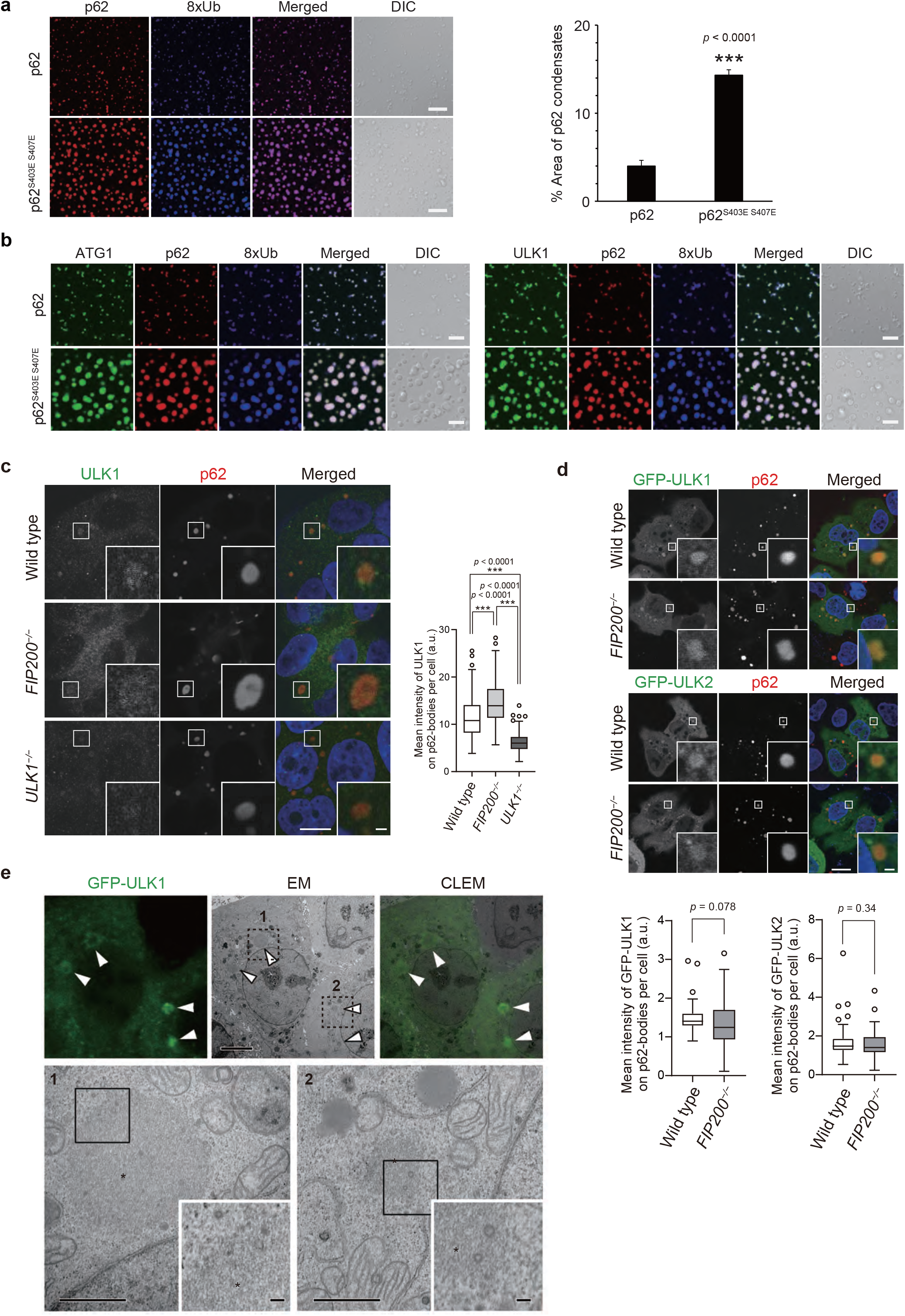
Localization of ULK1 on p62 bodies. **a** *In vitro* formation of p62-8xUb condensates. 10 μM mCherry-p62 wild-type or mCherry-p62^S403E S407E^, 10 μM SNAP-8xUb labelled with Alexa Fluor 649. Scale bars: 20 μm. The graph indicates the quantified area of p62 condensates formed by wild-type or mutant p62. Data are means ± s.d of wild-type p62 and p62^S403E, S407E^ condensates (*n* = 4). \*\*\**p* < 0.001 as determined by two-sided Welch’s t-test. **b** *In vitro* formation of p62-8xUb-Atg1 or ULK1 condensates. 10 μM mCherry-p62 wild-type or mCherry-p62^S403E, S407E^, 10 μM 8xUb labelled with Alexa Fluor 649 and 0.2 μM SNAP-Atg1 or SNAP-ULK1 labelled with Alexa Fluor 488 were mixed and observed by fluorescence microscopy. Scale bars: 10 μm. **C** Immunofluorescence microscopy. Wild-type, *FIP200*- or *ULK1*-knockout Huh-1 cells were immunostained with indicated antibodies. The mean fluorescence intensities of ULK1 on p62 bodies per cell were quantified in each genotype (*n* = 249 cells). Horizontal bars indicate medians, boxes the interquartile range (25th–75th percentiles) and whiskers 1.5× the interquartile range; outliers are plotted individually. Statistical analysis was performed by Šidák’s test after one-way ANOVA (\*\*\**p* < 0.001). Scale bars, 10 μm (main panels), 1 μm (inset panels). **d** Immunofluorescence microscopy. Wild-type and *FIP200*-knockout Huh-1 cells were transfected with GFP-ULK1 or GFP-ULK2 and immunostained with anti-p62 antibody. The mean fluorescence intensities of ULK1 on p62 bodies in each cell were quantified for each genotype (*n* = 79 cells). Horizontal bars indicate medians, boxes indicate interquartile range (25th–75th percentiles), and whiskers indicate 1.5× interquartile range; outliers are plotted individually. Statistical analysis was performed by Welch’s *t*-test. Scale bars, 10 μm (main panels), 1 μm (inset panels). **e** Correlative light and electron microscopy (CLEM) of Huh-1 cells expressing GFP-ULK1. Images of GFP-ULK1, corresponding electron micrograph (EM) images, and the merging of both (CLEM) are shown. Areas 1 and 2 are magnified in the bottom. Arrowheads indicate GFP-ULK1-positive p62 bodies. Scale bars, 5 μm (upper panel), 1 μm (lower panels), and 100 nm (insets of lower panels).

We next studied the localization of ULK1 in Huh-1 cells. Immunofluorescence analysis with an anti-ULK1 antibody showed the significant signal of ULK1 in p62 bodies, which was diminished by ULK1 depletion (Fig. 2c, Supplementary Fig. S2). Together with FIP200, ATG13, and ATG101, ULK1 forms an initiation kinase complex for autophagosome formation ^22^, and p62 interacts with FIP200 through the Claw domain ^13^, raising the possibility that the localization of ULK1 to p62 bodies is indirect and depends on the interaction of p62 with FIP200. To test this hypothesis, we developed *FIP200*-deficient Huh-1 cells (Supplementary Fig. S2). Remarkably, we observed ULK1 localization to p62 bodies even in these cells, and ULK1 signal intensity was significantly higher than in wild-type Huh-1 cells (Fig. 2c), probably due to increased ULK1 protein in the *FIP200*-knockout cells. Exogenously expressed green fluorescent protein (GFP)-tagged ULK1 and ULK2 also localized on p62 bodies regardless of the presence of FIP200 (Fig. 2d). Correlative light and electron microscopy with Huh-1 cells harboring GFP-ULK1 revealed that GFP-ULK1 localizes on round structures composed of filamentous assemblies; these structures were previously identified as p62 bodies ^15,23^ (Fig. 2e). Taken together, these data suggest that ULK1 localizes in p62 bodies through the direct ULK1-p62 interaction.

### Significance of ULK1 within p62 bodies

To investigate the significance of ULK1 and ULK2 within p62 bodies, we utilized MRT68921, which is the most potent inhibitor of ULK1 and ULK2, with IC_50_ values of 2.9 nM and 1.1 nM, respectively ^24^. As predicted, the treatment of Huh-1 cells with MRT68921 decreased not only the level of phosphorylated ATG13, but also those of the Ser349- and Ser403-phosphorylated p62 forms (Fig. 3a and Supplementary Fig. S3). Similar results were obtained with ULK-101, another inhibitor of ULK1 and ULK2 ^25^ (Supplementary Fig. S4).

**Fig. 3.**
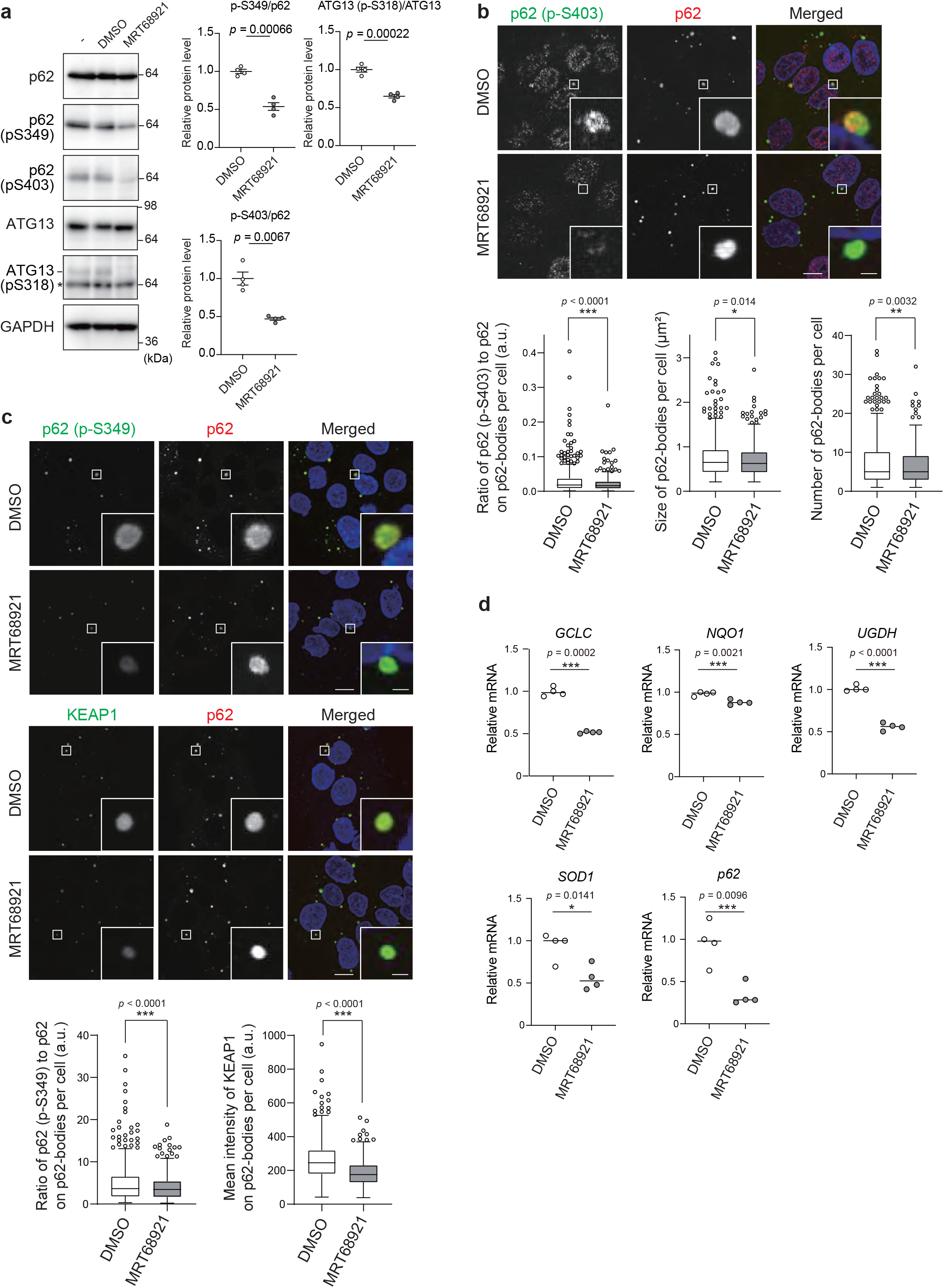
Significance of p62 phosphorylation by ULK1 and ULK2. **a** Immunoblot analysis. Huh-1 cells were treated with or without 2.5 μM MRT68921 for 6 h, and the cell lysates were subjected to immunoblot analysis with indicated antibodies. The asterisk indicates non-specific bands. Data shown are representative of three separate experiments. Bar graphs show the results of quantitative densitometric analysis of Ser349- or Ser403-phosphorylated p62 forms relative to total p62 (*n* = 3), and of Ser318-phosphorylated ATG13 relative to total ATG13 (*n* = 3). Data are means ± s.e. Statistical analysis was performed by Welch’s *t*-test. **b** Immunofluorescence microscopy. Huh-1 cells were treated with or without 2.5 μM MRT68921 for 6 h and immunostained with the indicated antibodies. The ratio of p62 (p-S403) to p62 on p62 bodies and the size and number of p62 bodies in each cell were quantified (*n* = 500 cells). Horizontal bars indicate medians, boxes indicate interquartile range (25-75th percentiles), and whiskers indicate 1.5× interquartile range; outliers are plotted individually. Statistical analysis was performed by Welch’s *t*-test (\**p* < 0.05, \*\**p* < 0.01, and \*\*\**p* < 0.001). Scale bars, 10 μm (main panels), 1 μm (inset panels). **c** Immunofluorescence microscopy. Huh-1 cells were treated with or without 2.5 μM MRT68921 for 6 h and immunostained with the indicated antibodies. The ratio of p62 (p-S349) to p62 and the signal intensity of KEAP1 on p62 bodies in each cell were quantified (*n* = 500 cells). Horizontal bars indicate medians, boxes indicate interquartile range (25th-75th percentiles), and whiskers indicate 1.5× interquartile range; outliers are plotted individually. Statistical analysis was performed by Welch’s *t*-test (\*\*\**p* < 0.001). Scale bars, 10 μm (main panels), 1 μm (inset panels). **d** Gene expression of NRF2 targets. Total RNAs were prepared from Huh-1 cells treated with or without 2.5 μM MRT68921 for 6 h. Values were normalized against the amount of mRNA in non-treated Huh-1 cells. qRT-PCR analyses were performed as technical replicates on each biological sample. Data are means ± s.e. \**p* < 0.05, and \*\*\**p* < 0.001 as determined by two-sided Welch’s *t*-test.

We observed extensive co-localization of the Ser403-phosphorylated form in p62 bodies (Fig. 3b). The signal intensity of Ser403-phosphorylated p62 in p62 bodies became weaker when Huh-1 cells were treated with MRT68921 (Fig. 3b). MRT68921 treatment slightly but significantly decreased both the size and number of p62 bodies (Fig. 3b). These results suggest that while ULK1 and ULK2 contribute to the LLPS of p62 through the phosphorylation of Ser403 of p62, the dephosphorylation of Ser403 within p62 bodies hardly have an effect on already formed p62 bodies.

Next, we tested whether the inhibition of ULK1 and ULK2 affected KEAP1-localization within p62 bodies. Huh-1 cells were cultured in the presence or absence of MRT68921 and immunostained with anti-p62 and anti-Ser349-phosphorylated p62-specific antibodies. The p62 bodies in Huh-1 cells contained the Ser349-phosphorylated form (Fig. 3c). Upon exposure to MRT68921, the signal intensities of phosphorylated p62 on p62 bodies markedly decreased (Fig. 3c). Double immunofluorescence analysis with anti-p62 and anti-KEAP1 antibodies showed extensive localization of KEAP1 in the p62 bodies (Fig. 3c). The signal intensity of KEAP1 in the bodies was significantly attenuated by treatment with MRT68921 (Fig. 3c), suggesting release of KEAP1 from the bodies to the cytoplasm and subsequent NRF2 inactivation. Indeed, gene expression of NRF2 targets such as glutamate-cysteine ligase catalytic subunit (*GCLC*), NAD(P)H quinone dehydrogenase 1 (*NQO1*), UDP-glucose 6-dehydrogenase (*UGDH*), superoxide dismutase 1 (*SOD1*), and *p62* itself was decreased by MRT68921 treatment (Fig. 3d). These data suggest that ULK1 and ULK2 in p62 bodies contribute to the activation of NRF2 by phosphorylating p62 at Ser349 and promoting sequestration of KEAP1 within p62 bodies.

### Dynamics of KEPA1 in S349-phosphorylated p62 bodies

In the next series of experiments, we sought to determine whether Ser349 phosphorylation of p62 affects KEAP1 dynamics in p62 bodies. To do this, we generated *p62 KEAP1* double-knockout Huh-1 cells (Supplementary Fig. S2) and expressed GFP-tagged wild-type p62, phosphomimetic p62^S349E 20^, phosphodefective p62^S349A 20^, or KEAP1 interaction-defective p62^T350A 19^ together with mCherry or mCherry-tagged KEAP1. The fluorescence analysis revealed that in the absence of mCherry-KEAP1, wild-type GFP-p62 and all GFP-p62 mutants formed round, liquid droplet-like structures (Fig. 4a). When we coexpressed wild-type GFP-p62 or the mutants with mCherry-KEAP1 in the double knockout Huh-1 cells, mCherry-KEAP1 co-localized well with GFP-p62-positive structures except those composed of GFP-p62^T350A^ (Fig. 4a). We measured the circularity of each GFP-p62-positive structure composed of wild-type p62 or the mutants in the presence or absence of mCherry-KEAP1. Circularity values close to 1 were associated with liquid droplets, while lower values correlated with gels or aggregates ^26,27^. In the absence of mCherry-KEAP1, all GFP-p62-positive structure values were close to 1 (Fig. 4b), suggesting that they were liquid droplets. Remarkably, the circularity of GFP-p62^S349E^ bodies but not others was significantly decreased when these bodies co-localized with mCherry-KEAP1 (Fig. 4b). Further, *in vitro* LLPS assay revealed that introduction of the S349E mutation in p62^S403E S407E^ resulted in the formation of amorphous aggregates rather than liquid droplets, but only in the presence of KEAP1 (Fig. 4c and Supplementary Fig. S5). These data suggest that strong binding of KEAP1 to p62 as a result of the S349E mutation changes the biophysical properties of p62 bodies.

**Fig. 4.**
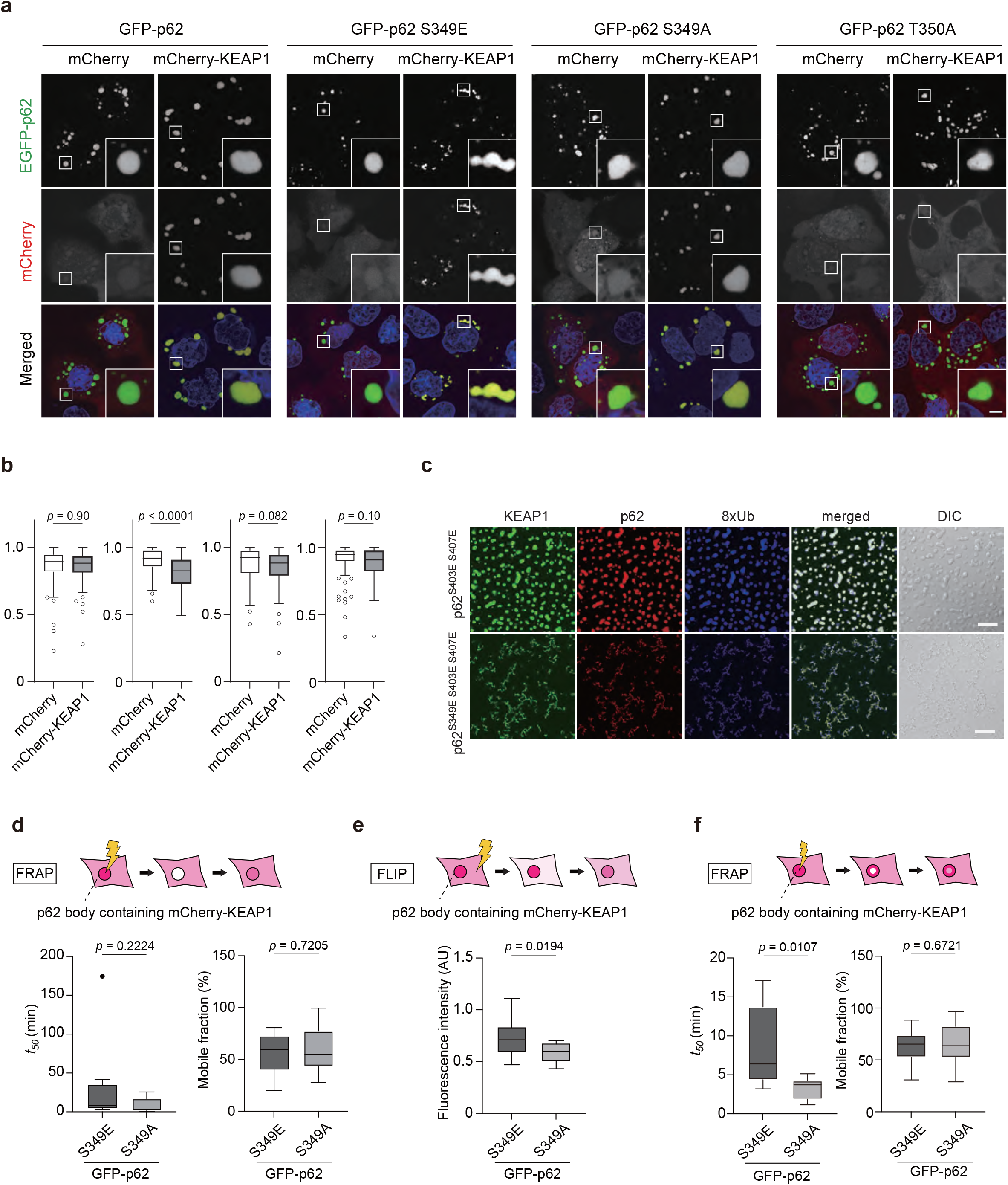
Dynamics of KEAP1 in p62 bodies. **a** Fluorescence microscopy. GFP-p62, GFP-p62^S349E^, GFP-p62^S349A^, or GFP-p62^T350A^ were co-transfected with mCherry or mCherry-KEAP1 into Huh1 *p62/KEAP1* double-knockout cells. Twenty-four hours after transfection, the fluorescence images were observed. **b** Circularity of p62 bodies. The circularity of p62 bodies in each cell was quantified (*n* = 150 cells). Horizontal bars indicate medians, boxes indicate interquartile range (25th–75th percentiles), and whiskers indicate 1.5× interquartile range; outliers are plotted individually. Statistical analysis was performed by two-sided Welch’s *t*-test. Scale bar: 2 μm. **c** *In vitro* formation of p62-KEAP1-8xUb condensates. 5 μM SNAP-KEAP1 labeled with SNAP-Surface Alexa Fluor 488 was premixed with 10 μM mCherry-p62 mutants before mixing with 10 μM SNAP(649)-8xUb. Scale bars: 20 μm. **d** FRAP analyses of mCherry-KEAP1 localized in p62 bodies comprised of GFP-p62^S349E^ or GFP-p62^S349A^. The half-time of recovery (*t50*) and mobile fraction (MF) of mCherry-KEAP1 was measured by FRAP of whole p62 bodies (*n* = 7). Data are means ± s.d. Statistical analysis was performed by two-sided Welch’s t-test. **e** FLIP analyses of mCherry-KEAP1 localized in p62 bodies comprised of GFP-p62^S349E^ or GFP-p62^S349A^. The fluorescence loss of mCherry-KEAP1 in GFP-p62 bodies (*n* = 14) was measured 30 min after photobleaching over a large area of cells. **f** FRAP analyses of mCherry-KEAP1 localized in p62 bodies comprised of GFP-p62^S349E^ or GFP-p62^S349A^. *t50* and MF of mCherry-KEAP1 were measured by FRAP of the central portions of p62 bodies (*n* = 10). Data are means ± s.d. Statistical analysis was performed by two-sided Welch’s t-test.

We hypothesized that once mCherry-KEAP1 was incorporated into S349-phosphorylated p62 bodies, KEAP1 efflux from the bodies would be significantly reduced due to the close interaction between the phosphorylated p62 and KEAP1. To prove this, we used fluorescence recovery after photobleaching (FRAP) and fluorescence loss in photobleaching (FLIP) ^28^ to evaluate KEAP1 influx and efflux into p62 bodies consisting of GFP-p62^S349E^ or p62^S349A^. To measure the influx of mCherry-KEAP1 into GFP-p62 bodies, the whole fluorescence of mCherry-KEAP1 in GFP-p62^S349E^ or p62^S349A^ bodies was photobleached, and the fluorescence recovery was measured. It is common to characterize molecular dynamics in FRAP experiments by the half-time of recovery (*t50*) and the mobile fraction ^29^. Based on these parameters, the influx of mCherry-KEAP1 from the surrounding environment was comparable between p62^S349E^ and p62^S349A^ bodies (Fig. 4d and Supplementary Movies S5 and S6). Next, to examine the efflux of mCherry-KEAP1 from GFP-p62 bodies, we carried out FLIP analysis. When approximately 80% of the cellular region is photobleached, the fluorescent signal in the cytoplasm is transiently reduced, followed by a gradual recovery due to influx from the non-bleached region. If there is an outflow of mCherry-KEAP1 from p62 bodies, the fluorescence intensity of mCherry-KEAP1 within p62 bodies in the non-bleached area should decrease after photobleaching. While the signal intensity of mCherry-KEAP1 in p62^S349A^ bodies decreased about to 58.2 ± 0.05% of the baseline value at 30 min after the photobleaching, it remained higher (72.0 ± 0.07%) in the case of p62^S349E^ bodies (Fig. 4e and Supplementary Movies S7 and S8). Finally, to investigate the inner fluidity of mCherry-KEAP1 in GFP-p62 bodies, we measured the fluorescence recovery of mCherry-KEAP1 after photobleaching of the central portion of GFP-p62 bodies. The *t50* of mCherry-KEAP1 in p62^S349E^ bodies was 8.51 ± 4.83 min, which was much slower than that seen with p62^S349A^ bodies (3.32 ± 1.23 min) (Fig. 4f and Supplementary Movie S9 and S10). Taken together, these results suggest that Ser349 phosphorylation of p62 results in the retention of KEAP1 in p62 bodies, and when this sequestration is prolonged, the inner fluidity of these bodies is decreased.

### Physiological significance of Ser349 phosphorylation of p62 in mice

To clarify the physiological role of Ser349 phosphorylation of p62 *in vivo*, we generated knock-in mice that expressed p62 in which Ser351 (corresponding to human Ser349) was replaced by Glu (*p62*^*S351E/+*^ mice) or Ala (*p62*^*S351A/+*^ mice). Initially, we tried to use the CRISPR/Cas9 system to generate both knock-in mice, but could obtain only *p62*^*S351A/+*^ mice. We therefore attempted to use prime editing, a recently developed system, to generate *p62*^*S351E/+*^ mice using mouse embryonic stem cells (mES cells) ^30^. Even with this method, however, we were unable to obtain chimeric mice with high chimerism. We did succeed in generating a male chimeric mouse with low chimerism and germline transmission. *In vitro* fertilization using sperm from the chimeric mouse was performed to obtain a sufficient number of heterozygotes for the experiments. The resulting *p62*^*S351E/+*^ mice showed severe growth retardation and mild hepatomegaly at P12 and P15 (Fig. 5a-c), which is probably why knock-in mice could not be obtained by the above method. RNAseq analysis of wild-type and *p62*^*S351E/+*^ mouse livers demonstrated increased gene expression of NRF2 targets (Fig. 5d). Consistent with these results, real-time PCR analysis showed that the gene expression of NRF2 targets such as *glutathione S-transferase Mu 1* (*Gstm1*), *Nqo1, Ugdh*, and *p62* was much higher in the liver of *p62*^*S351E/+*^ mice than in wild-type mice (Fig. 5e). We also found that the levels of GSTM1, NQO1, and UGDH, and the nuclear level of NRF2, were markedly higher in *p62*^*S351E/+*^ mice than in wild-type mice (Fig. 5f). These results indicate that the expression of *p62*^*S351E*^ at half the level of endogenous *p62* is sufficient for KEAP1 inactivation and subsequent NRF2 activation *in vivo*. Anatomical analysis revealed that the forestomach wall of *p62*^*S351E/+*^ mice was obviously thickened compared with that of wild-type mice (Supplementary Fig. S6). Hematoxylin and eosin (HE) staining indicated that the esophagus and forestomach of *p62*^*S351E/+*^ mice had a remarkably thicker stratum corneum than those of wild-type mice, but such differences were not observed in the skin (Fig. 5g). In the forestomach, the epithelial layers below the stratum corneum were slightly thicker in *p62*^*S351E/+*^ mice than in wild-type mice, but this was not evident in the esophagus. Immunohistochemistry (IHC) revealed that in the epithelia of the esophagus and forestomach, the intensity of NQO1 was higher in *p62*^*S351E/+*^ mice than in wild-type mice (Fig. 5h). The staining of the cell proliferation marker Ki-67 showed no significant difference between mutant and wild-type organs (data not shown). Serum data from *p62*^*S351E/+*^ mice indicated a slightly but significantly elevated aspartate aminotransferase level, signs of malnutrition (low blood glucose and high cholesterol), and signs of dehydration (increased blood urea nitrogen and creatinine) (Fig. 5i). These results strongly suggested that *p62*^*S351E/+*^ mice are a phenocopy of *Keap1*-deficient mice ^31^, and that impaired nutritional intake due to hyperkeratosis in the esophagus and forestomach could be the primary cause of the severe phenotype of *p62*^*S351E/+*^ mice. In striking contrast to these mice, *p62*^*S351A/+*^ and *p62*^*S351A/S351A*^ mice were fertile and showed no obvious growth retardation (Supplementary Fig. S7a-c). Morphological and biochemical analyses indicated no differences in phenotypes (including NRF2 activation) between wild-type and *p62*^*S351A/S351A*^ mice (Supplementary Fig. S7d-g), implying that Ser351 phosphorylation of p62 is unnecessary for mouse development and survival. Taken together, these results indicate that Ser351 phosphorylation of p62 is physiologically important for the control of NRF2 activity *in vivo*, most likely by regulating the redox-independent KEAP1-NRF2 pathway.

**Fig. 5.**
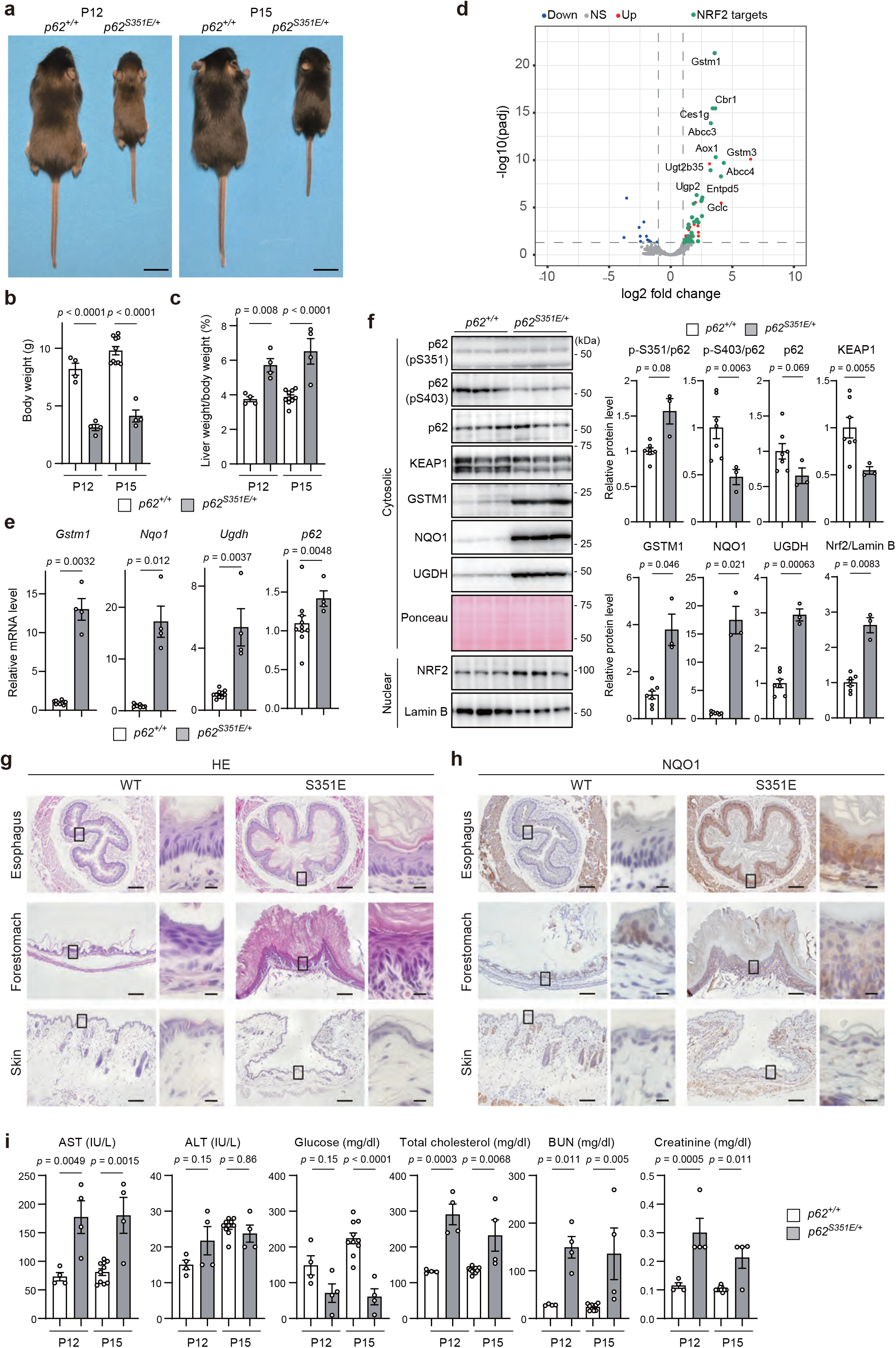
Physiological significance of p62 phosphorylation at S351 in mice. **a** External appearance of *p62*^*+/+*^ and *p62*^*S351E/+*^ mice at postnatal days (P) 12 and 15. **b, c** Body weight (g) (**b**) and liver weight (% of body weight) (**c**) of *p62*^*+/+*^ (*n* = 4 at P12, *n* = 10 at P15) and *p62*^*S351E/+*^ mice (*n* = 4). Data are means ± s.e. Statistical analysis was performed by Tukey’s test after one-way ANOVA. **d** RNA-seq analysis of livers of *p62*^*+/+*^ and *p62*^*S351E/+*^ mice at P19 (*n* = 3). Volcano plots of differentially expressed genes in *p62*^*+/+*^ versus *p62*^*S351E/+*^ mice (red points: FDR < 0.05 and log2FC > 1, blue points: FDR < 0.05 and log2FC < -1). Green points are the targets of transcription factor NRF2. **e** Gene expression of NRF2 targets. Total RNAs were prepared from mouse livers of *p62*^*+/+*^ (*n* = 10) and *p62*^*S351E/+*^ (*n* = 4) mice at P15. Data are means ± s.e. Statistical analysis was performed by Welch’s *t*-test. **f** Immunoblot analysis of *p62*^*+/+*^ (*n* = 7) and *p62*^*S351E/+*^ (*n* = 3) mice at P19. Liver homogenates were subjected to immunoblot analysis with the indicated antibodies. Bar graphs show the results of quantitative densitometric analysis. Data are means ± s.e. Statistical analysis was performed by Welch’s *t*-test. **g, h** Hematoxylin and eosin (HE) staining (**g**) and immunohistochemical analysis of NQO1 (**h**) of livers from *p62*^*+/+*^ and *p62*^*S351E/+*^ mice at P19. Scale bars, XX μm. **i** Serum levels of aspartate aminotransferase (AST), alanine aminotransferase (ALT), glucose, total cholesterol, blood urea nitrogen (BUN), and creatinine from *p62*^*+/+*^ (*n* = 4 at P12, *n* = 10 at P15) and *p62*^*S351E/+*^ (*n* = 4) mice at P12 and P15 were measured. IU/l, international units/liter. Data are means ± s.e. Statistical analysis was performed by Tukey’s test after one-way ANOVA.

## Discussion

The ULK1 kinase complex functions as the most upstream factor for autophagosome formation ^22,32^. In *Saccharomyces cerevisiae*, nutrient deprivation causes phosphatase-induced dephosphorylation of Atg13, which results in the formation of higher-order structures of the Atg1 kinase complex (the ULK1 kinase complex in mammals) and subsequently the formation of a liquid-droplet, pre-autophagosomal structure ^33,34^. Nutrient starvation activates the ULK1 kinase complex, which phosphorylates ATG proteins, including Beclin 1 ^35^, ATG14 ^36,37^, and ATG9 ^38^, and contributes to the initiation of autophagosome formation ^22,32^. On the other hand, ULK1 phosphorylates substrates that are not directly involved in autophagosome formation, such as SEC16A ^39^, glycolytic enzymes ^40^, and STING ^41^. p62 is also in this category, and it is closely involved in the LLPS, the p62 body formation ^42^. In this study, we showed that ULK1 localized to p62 bodies (Fig. 2) and phosphorylated Ser349 of p62 (Fig. 1), the latter of which is required for KEAP1 localization and retention to p62 bodies and subsequent NRF2 activation (Figs. 3 and 4). Thus, ULK1 modulates the formation and degradation of p62 bodies and also plays a role in the antioxidative-stress response.

When does ULK1 phosphorylate p62? Since ULK1 was localized in p62 bodies both *in vitro* and *in vivo* (Fig. 2), it is plausible that ULK1 phosphorylates Ser349 of p62 located in p62 bodies. Indeed, we hardly observed phosphorylation of the p62^K7A D69A^ mutant, which is not capable of LLPS (data not shown). How does ULK1 recognize p62 bodies? Our HS-AFM showed that the p62 homodimer directly bound to ULK1 via dynamic IDR-IDR and IDR-globular domain interactions (Fig. 1f). It is known that IDRs specifically interact with multiple target molecules through a binding mode called “coupled folding and binding” ^43^. This binding mode may facilitate conformational changes and phosphorylation of large numbers of p62 molecules in p62 bodies, because substitutions between IDRs that bind to target molecules occur very rapidly ^44^. With the phosphomimetic p62 mutant p62^S349E^, influx of KEAP1 into p62 bodies predominated over efflux (Fig. 4d and e). The coexpression of p62^S349E^ and KEAP1 in *p62 KEAP1* double-knockout Huh-1 cells reduced the circularity of p62 bodies compared to the expression of p62^S349E^ alone (Fig. 4a and b), and also decreased the fluidity of KEAP1 molecules within p62 bodies (Fig. 4f). When the sequestration of KEAP1 within p62 bodies surpasses a certain threshold level, these bodies convert from liquid-like to gel-like droplets. What does this mean? Since autophagy is known to target gel-like rather than liquid-like droplets ^15,45,46^, it is possible that incorporation of a certain number of KEAP1 molecules into p62 bodies changes them into gel-like aggregates and enhances autophagic degradation. This is consistent with two findings: adding KEAP1 to p62 condensates consisting of p62^S349E^ resulted in amorphous aggregates *in vitro* (Fig. 4c), and *p62*^*S351E/+*^ mice had reduced levels of not only Ser403-phosphorylated p62 representing p62 bodies but also KEAP1 (Fig. 5f). Once p62 bodies are degraded by autophagy, the interaction between Ser349-phosphorylated p62 and KEAP1 should be suppressed because Ser349 phosphorylation of p62 occurs mainly in p62 bodies. As a result, KEAP1 remains in the cytoplasm, and NRF2 is degraded. In other words, the retention of KEAP1 in p62 bodies above a threshold level is thought to suppress NRF2 activation in a feedback regulation process.

*Nrf2*^*-/-*^ mice grow normally and are fertile ^47^, though they are susceptible to oxidative stress and reactive electrophiles ^17^. In addition, they show tooth decolorization due to defective iron transport in the enamel ^48^, which makes them easily distinguishable from wild-type and heterozygous mice. *p62*^*S351A/S351A*^, the phosphodefective *p62* knock-in mice in which p62-mediated NRF2 activation should be impaired, were also fertile and did not differ from wild-type mice for 1 year, at least under specific pathogen-free conditions (Supplementary Fig. S7). However, their incisors were brownish-yellow, in contrast to the greyish white incisors in *Nrf2*-deficient mice (data not shown), indicating that *p62*^*S351A/S351A*^ mice retain NRF2 activity. p62-mediated NRF2 activation was not required for development or survival, at least under steady-state conditions. The redox-independent antioxidative stress response might have an anti-aging effect since p62 bodies are thought to increase when the activities of both autophagy-lysosomes and the ubiquitin-proteasome system decrease with aging. Further research is needed to determine whether the redox-independent stress response is activated with aging, and if so, which tissues are affected.

In sharp contrast to *p62*^*S351A/S351A*^ mice, the phosphomimetic *p62* knock-in mice *p62*^*S351E/+*^, in which p62-mediated NRF2 activation is persistently activated, showed severe phenotypes. The mice had impaired nutritional intake due to hyperkeratosis in the esophagus and forestomach, which led to malnutrition and dehydration (Fig. 5). This represented almost a phenocopy of *Keap1*-deficient mice, which exhibit hyperactivation of NRF2 and show hyperkeratosis in the esophagus and forestomach ^31^. One difference is that *p62*^*S351E/+*^ mice do not develop skin hyperkeratosis, which is observed in *Keap1*-knockout mice ^31^. Why does constant activation of NRF2, which has an inherently cytoprotective role, cause a severe phenotype? The esophagus and stomach are exposed to a variety of toxic foods and drinks. This may cause wounds that require NRF2 for healing, and a redox-independent p62-mediated pathway may be at work. Meanwhile, the healing of skin wounds may be mediated by redox-dependent NRF2 activation ^49^. It is plausible that transient activation of NRF2 in response to toxicity, whether redox-dependent or redox-independent, is important for biological defense, and that persistent activation leads to excessive defense responses (*e*.*g*., excessive keratinization). In the case of redox-independent stress response, the transient activation would be regulated by phosphorylation, dephosphorylation, and autophagic degradation of p62 body.

In conclusion, we showed for first time that the redox-independent NRF2 activation pathway, which is mediated by p62 bodies and their phosphorylation, is physiologically important.

## Methods Cell culture

Huh-1 cells (JCRB0199, NIBIOHN) were cultured in Dulbecco’s modified Eagle’s medium containing 10% fetal bovine serum, 5 U/mL penicillin, and 50 μg/mL streptomycin. For overexpression experiments, Huh-1 cells were transfected using Lipofectamine 3000 (L3000015, Thermo Fisher Scientific, Waltham, MA, USA). Huh-1 cells were authenticated using the STR profile and tested for mycoplasma contamination.

### Mice

To generate *p62*^*S351A*^ knock-in mice, CRISPR RNA (crRNA) was designed to recognize the target site (5’-ACTGGAGTTCACCTGTAGAT-3’). The synthetic crRNAs (Alt-R CRISPR-Cas9 crRNA), trans-activating CRISPR RNA (tracrRNA) (Alt-R CRISPR-Cas9 tracrRNA), and Cas9 protein (Alt-R S.p. Cas9 Nuclease V3) were purchased from Integrated DNA Technologies, Inc. (IDT; Coralville, IA, USA). The 120-mer single-stranded oligodeoxynucleotide (ssODN) carrying the intended base substitutions (TCT to GCC) was synthesized by Eurofins Genomics K.K. (Tokyo, Japan). The CRISPR/Cas9 solution was prepared as previously described ^50^, with minor modifications. Briefly, lyophilized crRNAs and tracrRNA were resuspended in nuclease-free duplex buffer (IDT) to a concentration of 240 μM. Equal volumes of crRNA and tracrRNA were combined, heated at 95°C for 5 min, and then placed in room temperature (RT) for about 10 min to allow formation of crRNA-tracrRNA duplex. Lyophilized ssODN was resuspended in nuclease-free water to a concentration of 4 mg/mL. crRNA-tracrRNA duplex was mixed with Cas9 protein to form a ribonucleoprotein complex, and then mixed with ssODN in Opti-MEM (Thermo Fisher Scientific). The final concentrations of Cas9 protein, crRNA-tracrRNA duplex, and ssODN were 1 μg/mL, 30 μM, and 1mg/mL, respectively. To induce CRISPR/Cas9-mediated mutation, we applied a method called improved genome editing via oviductal nucleic acid delivery (i-GONAD) ^50^. Approximately 1.5 mL of CRISPR/Cas9 solution was injected into the oviductal lumens of female C57BL/6N mice on day 0.7 of pregnancy. Immediately after the injection, the oviduct regions were grasped with a tweezer-type electrode (catalog no. CUY652-3; Nepa Gene Co., Ltd., Chiba, Japan) and then electroporated using the NEPA21 square-wave pulse generator (Nepa Gene). The electroporation parameters used were previously described ^50^. Pregnant female mice were allowed to deliver their pups. Biopsies of pup tails were performed for genomic DNA isolation, and mutations were validated by sequencing of PCR products amplified from genomic DNAs.

To generate *p62*^*S351E*^ knock-in mice using mouse embryonic stem (mES) cells, we applied the recently developed prime editing system ^30^. Details of the methods for establishment of edited mES cells will be published elsewhere (Manabu Abe, et al). Briefly, we designed prime-editing guide RNA (pegRNA) containing the following sequences: spacer sequence, 5’-GACUGGAGUUCACCUGUAGA-3’; reverse transcription template, 5’-GUGGACCCAGAG-3’; primer-binding site, 5’-ACAGGUGAACUCC-3’. The CAG promoter-driven prime editor 2 (PE2) and the U6 promoter-driven pegRNA expression vectors were originally constructed using pCMV-PE2-P2A-GFP (#132776, Addgene) and the hU6-sgRNA plasmid ^51^. These vectors were co-transfected into RENKA4, a C57BL/6N-derived mES cell line, using Lipofectamine 3000 (Thermo Fisher Scientific). Knock-in mutations in transfected mES clones were validated by sequencing of PCR products amplified from genomic DNAs. Culture of mES cells and generation of chimeric mice were carried out as previously described ^52^.

### Purification of recombinant protein

mCherry-p62, mCherry-p62 mutants, SNAP-KEAP1, and SNAP-8xUb were prepared as described previously ^15^. The gene encoding 8xUb was purchased from GenScript, New Jersey,USA. Purified proteins were stored at −80 °C until use.

Recombinant SNAP-Atg1 proteins were prepared as described previously ^34^. To construct the expression plasmid encoding TwinStrep-CS-SNAP-ULK1-His6 (CS; HRV 3C protease recognition site), the genes and pCAG-neo vector (FUJIFILM Wako Pure Chemical Corporation) were amplified by PCR. The PCR fragments were assembled using NEBuilder HiFi DNA Assembly Master Mix (New England BioLabs (NEB), Ipswich, MA). The construct was subjected to sequencing analysis to confirm its identity and transfected into Expi293 GnTI− cells using Screen *F*ect™ UP-293 (FUJIFILM Wako Pure Chemical Corporation). The cells were collected after 5 days and sonicated in lysis buffer (50 mM Tris-HCl [pH 8.0], 300 mM NaCl, 2 mM MgCl_2_, 1% Triton X-100, 10% glycerol, 1 mM TCEP, 1x protease inhibitor cocktail (Nacalai Tesque, Kyoto, Japan)) on ice and centrifuged at 16,000 *g* for 40 min at 4 °C. The supernatant was purified with a Strep-Tactin®XT resin column (IBA Lifesciences Göttingen, Germany). The protein was eluted with 20 mM Tris-HCl (pH 8.0), 500 mM NaCl, 50 mM biotin and concentrated using Vivaspin 500 (Cytiva, Massachusetts, USA).

### HS-AFM

The procedure for HS-AFM observation was described previously ^53^. HS-AFM images were acquired in tapping mode using a sample-scanning HS-AFM instrument (MS-NEX, Research Institute of Biomolecule Metrology Co., Ltd., Ibaraki, Japan). We used cantilevers measuring ∼7 μm long, ∼2 μm wide, and ∼0.08 μm thick with electron beam-deposited (EBD) tips (tip radius < 10 nm) (USC-F1.2-k0.15, NanoWorld, Neuchâtel, Switzerland). Their resonant frequency and spring constant were 1.2 MHz in air and 0.15 N/m, respectively. Imaging conditions were as follows: scan size, 120 × 120 nm^2^ (Fig. 2f, Fig. S3a, b) or 150 × 150 nm^2^ (Fig. S3c); pixel size, 100 × 100 pixels (Fig. 2f, Fig. S3a-c); imaging rate, 8.33 frames/sec (Fig. 2f, Fig. S3a) or 6.67 frames/s (Fig. S3b,c). Imaging was performed at 23 °C. HS-AFM images were viewed and analyzed using the Kodec4.4.7.39 ^54^ and ImageJ software systems.

### Sample preparation for HS-AFM imaging

For imaging of SNAP-ULK1 and p62_268–440, SNAP-ULK1 (50 nM) or p62_268–440 (10 nM) was deposited onto freshly cleaved mica glued to the top of a glass stage (diameter, 1.5 mm; height, 2 mm). After incubation for 3–5 min, the mica was rinsed and immersed in the liquid cell containing ∼90 μL of imaging buffer A (20 mM NaCl, 20 mM HEPES-NaOH [pH 7.5], 1 mM MgCl_2_, 0.1 mM ATP) or imaging buffer B (20 mM NaCl, 20 mM HEPES-NaOH [pH 7.5], 1 mM MgCl_2_), respectively. For imaging of p62_268–440 with SNAP-ULK1 or SNAP-ATG1, 25 nM p62_268–440 and 50 nM SNAP-ULK1 or 10 nM p62_268–440 and 5 nM SNAP-ATG1 in imaging buffer A were mixed in a 0.5-mL tube. The mixed protein solution was deposited onto freshly cleaved mica and incubated for 3–5 min. After rinsing with imaging buffer A, the mica was immersed in ∼90 μL of imaging buffer A.

### *In vitro* kinase assay

Purified p62 (268–440aa), p62 (320–440aa) or mCherry-p62 was incubated with purified SNAP-Atg1 or SNAP-ULK1 in kinase buffer (20 mM Tris-HCl [pH 7.5], 150 mM NaCl, 0.5 mM MgCl_2_, 0.1 mM DTT) containing 200 μM of ATP per reaction for 30 min at 37 °C. The reaction was terminated by adding LDS sample buffer (NP0007, Thermo Fisher Scientific) and subjected to SDS-PAGE followed by immunoblot analyses with p62 Ser349-^20^, Ser403-(GTX128171, GeneTex), and Ser407-^8^ specific antibodies.

### *In vitro* LLPS assay

For the *in vitro* LLPS assay, fluorescence observation was performed on glass-bottom dishes (MatTek, Massachusetts, USA) coated with 0.3% (w/v) bovine serum albumin using an FV3000RS confocal laser-scanning microscope (Olympus, Tokyo, Japan). Lasers with wavelengths of 488, 561, and 640 nm were used for excitation of Alexa Fluor 488, mCherry, and Alexa Fluor 649, respectively. To observe p62-8xUb condensates, mCherry-p62 wild-type (WT) or mCherry-p62 mutant was mixed with SNAP(649)-8xUb (SNAP-8xUb labeled with SNAP-Surface 649). Each final protein concentration was 10 μM. To observe p62-8xUb condensates in the presence of ATG1 or ULK1, SNAP(488)-ATG1 or SNAP(488)-ULK1 (SNAP-ATG1 / ULK1 labeled with SNAP-Surface Alexa Fluor 488) was premixed with mCherry-p62WT / p62S403_S407E before mixing with SNAP(649)-8xUb. The final concentrations of SNAP(488)-Atg1 / ULK1, SNAP(649)-8xUb, and mCherry-p62WT / p62S403E_S407E were 0.2 μM, 10 μM, and 10 μM, respectively. To observe p62-8xUb condensates in the presence of KEAP1, SNAP(488)-KEAP1 (SNAP-KEAP1 labeled with SNAP-Surface Alexa Fluor 488) was premixed with mCherry-p62 mutant before mixing with SNAP(649)-8xUb. The final concentrations of SNAP(488)-KEAP1, SNAP(649)-8xUb, and mCherry-p62 mutants were 5 μM, 10 μM, and 10 μM, respectively. The buffer solutions used for the *in vitro* LLPS assay were 200 mM NaCl, 20 mM HEPES-NaOH (pH 7.5), 10% glycerol, and 1 mM TCEP. Each mixed solution was incubated for 50 min at ∼23 °C before imaging.

The % area of p62 condensates (the sum of the areas of p62 condensates divided by the total area (106.7 μm × 106.7 μm)) (Fig. 2a) was calculated using the “analyze particles” tool of ImageJ software. Statistical analysis was performed by Welch’s *t*-test.

### Immunoblot analysis

Cells were lysed in ice-cold TNE buffer (50 mM Tris-HCl [pH 7.5], 150 mM NaCl, 1 mM EDTA) containing 1% Triton X-100 and cOmplete EDTA-free protease inhibitor cocktail (5056489001, Roche). After centrifugation twice at 15,000 *g* for 10 min, the supernatant was collected as the cell lysates. Protein concentrations were determined by bicinchoninic acid (BCA) protein assay (23225, Thermo Fisher Scientific). The lysate was boiled in LDS sample buffer, and the samples were separated by SDS-PAGE and then transferred to polyvinylidene difluoride membranes. For gene knockdown, cells were transfected with

siGENOME siRNA targeting *ULK1* (D-005049-01-0005; 5’-

CCUAAAACGUGUCUUAUUU-3’, D-005049-02-0005; 5’-

ACUUGUAGGUGUUUAAGAA-3’, D-005049-03-0005; 5’-

GGUUAGCCCUGCCUGAAUC-3’, Horizon Discovery, United Kingdom) and *ULK2* (MQ-005396-01-0002; mixtures of sequences 5’-UAAAGGAACUUCAGCAUGA-3’, 5’-GUGGAGACCUCGCAGAUUA-3’, 5’-GAAGAACAGUCGAAAGAUUA-3’, 5’-GCAGACGUGCUUCAAAUGA-3’, Horizon Discovery) or a non-targeting control siRNA (siGENOME Non-Targeting siRNA; mixtures of sequences 5’-UAGCGACUAAACACAUCAA-3’, 5’-UAAGGCUAUGAAGAGAUAC-3’, 5’-AUGUAUUGGCCUGUAUUAG-3’, and 5’-AUGAACGUGAAUUGCUCAA-3’, Horizon Discovery) using DharmaFECT1 (T-2002, Horizon Discovery) and lysed 96 h after transfection in TNE buffer containing 1% SDS. Antibodies against p62 (610832, BD Biosciences, New Jersey, USA), Ser403-phosphorylated p62 (GTX128171, GeneTex California, USA), Ser349-phosphorylated p62 ^20^, Ser405-phosphorylated p62 ^8^, ULK1 (8054, Cell Signaling Technology, Massachusetts, USA), ULK2 (A15244, ABclonal, TE Huissen, Netherlands), FIP200 (17250-1-AP, Proteintech Group, Illinois, USA), and NRF2 (H-300; Santa Cruz Biotechnology, California, USA) were used as primary antibodies. Blots were then incubated with horseradish peroxidase-conjugated secondary antibody (Goat Anti-Mouse IgG (H + L), 115-035-166, Goat Anti-Rabbit IgG (H + L) 111-035-144, and Goat Anti-Guinea Pig IgG (H + L), all from Jackson ImmunoResearch, Pennsylvania, USA) and visualized by chemiluminescence.

### Immunofluorescence analysis

Huh-1 cells on coverslips were washed with PBS and fixed with 4% paraformaldehyde (PFA) for 15 min at RT, permeabilized with 0.1% Triton X-100 in PBS for 5 min, and blocked with 0.1% (w/v) gelatin (G9391, Sigma-Aldrich, Darmstadt, Germany) in PBS for 20 min. Then, cells were incubated with primary antibodies in the blocking buffer for 1 h, washed with PBS, and incubated with secondary antibodies for 1 h. Antibodies against p62 (610832, BD Biosciences), S403-phosphorylated p62 (GTX128171, GeneTex), S349-phosphorylated p62 ^20^, KEAP1 (10503-2-AP, Proteintech), and ULK1 (8054, Cell Signaling Technology) were used as primary antibodies. Goat anti-Mouse IgG (H + L) Highly Cross-Adsorbed Secondary Antibody, Alexa Fluor 647 (A21236, Thermo Fisher Scientific) and Goat anti-Rabbit IgG (H + L) Cross-Adsorbed Secondary Antibody, Alexa Fluor 488 (A11008, Thermo Fisher Scientific) were used as secondary antibodies. Nuclei were stained with Hoechst 33342 (62249, Thermo Fisher Scientific). Cells were imaged using the FV1000 confocal laser-scanning microscope with FV10-ASW 04.01 (Olympus) and a UPlanSApo ×60 NA 1.40 oil objective lens. Contrast and brightness of images were adjusted using Photoshop 2021v25.0 (Adobe, California, USA). The number, size, and circularity of p62-positive punctae in each cell and the mean fluorescence intensity of each signal on p62-positive punctae were quantified using a Benchtop High-Content Analysis System (CQ1, Yokogawa Electric Corp., Tokyo, Japan) and CellPathfinder software (Yokogawa Electric Corp.).

### Correlative light and electron microscopic analysis

Huh-1 cells on coverslips etched with 150-μm grids (CS01885, Matsunami Glass Ind. Osaka, Japan) were transfected with pMRX-IP-GFP-ULK1 or pMRX-IP-GFP-ULK2. After 24 h, cells were fixed with 2% PFA–0.1% glutaraldehyde (GA) in 0.1 M PB (pH 7.4). Then, phase-contrast and fluorescence images were obtained using a confocal microscope (FV1000). After image acquisition, the cells were fixed again with 2% PFA and 2% GA in 0.1 M PB (pH 7.4), processed according to the reduced-osmium method ^55^, and embedded in Epon812. Areas containing cells of interest were trimmed, cut as serial 80-nm sections, and observed using an electron microscope (EM; JEM1400; JEOL). Light microscopy and EM images were aligned according to three p62 bodies using Photoshop CS6 (Adobe).

### Histological analyses

Mouse livers were excised, cut into small pieces, and fixed by immersion in 4% PFA–4% sucrose in 0.1 M PB, pH 7.4. After rinsing, they were embedded in paraffin for immunostaining. Paraffin sections of 3-μm thickness were prepared and processed for HE staining or IHC. For IHC, antigen retrieval was performed for 20 min at 98 °C using a microwave processor (MI-77, AZUMAYA, Japan) in 1% immunosaver (Nissin EM, Japan). Sections were blocked and incubated for 2 days at 4 °C with the following primary antibodies: rabbit polyclonal antibody against NQO1 (Abcam), followed by N-Histofine simple stain mouse MAX PO kit (NICHIREI BIOSCIENCES, Tokyo, Japan) using 3,39-diaminobenzidine. Images of the stained specimens were acquired with a microscope (BX51, Olympus) equipped with a cooled CCD camera system (DP-71, Olympus).

### FRAP and FLIP assays

*p62*/*KEAP1* double-knockout Huh-1 cells expressing GFP-p62^S349E^ or GFP-p62^S349A^ in a doxycycline treatment-dependent manner were generated using a reverse tet-regulated retroviral vector, as previously reported. To induce the expression of GFP-p62^S349E^ or GFP-p62^S349A^, the cells were treated with 50 ng/mL of doxycycline (Dox, Sigma-Aldrich) for 24 h. Thereafter, mCherry-KEAP1 was transfected with Lipofectamine 3000 (Thermo Fisher Scientific) and cultured for 24 h. In FRAP assays, GFP-p62 bodies that were positive for mCherry-KEAP1 were bleached using a laser intensity of 10% at 488 nm, and then the fluorescence recovery of mCherry was recorded. In FLIP assays, almost 80% of the total cell area was set as the region of interest for photobleaching (excitation output level: 10% at 561 nm; iterations: 3–5) using FV31S-SW software (Olympus), and then the fluorescence loss of mCherry-KEAP1 in GFP-p62^S349E^ or GFP-p62^S349A^ bodies was recorded. The fluorescence intensity of mCherry-KEAP1 within the p62 bodies in the photobleaching area recovered to 32% at 15 min after photobleaching and reached equilibrium. Therefore, the fluorescence intensity of mCherry-KEAP1 within the p62 bodies in the non-photobleaching area was measured at 30 min after fluorescence loss. Olympus FV31S-SW software (version: 2.4.1.198) and cellSens Dimension Desktop 3.2 (Build 23706) was used for image collection and analysis. The mobile fraction was calculated from 10 measurements by the following equation: Mf = (F∞−F0)/(Fi−F0), where Mf is the mobile fraction, F∞ is the fluorescence intensity after full recovery (plateau), Fi is the initial fluorescence intensity prior to bleaching, and F0 is the fluorescence intensity immediately after bleaching. The half-time (*t50*) of fluorescence recovery was calculated from 10 measurements by curve fitting using the one-phase decay model of GraphPad PRISM 9 (GraphPad Software, California, USA).

### RNA sequencing (RNA-seq)

Total RNA from livers of *p62*^*+/+*^ and *p62*^*S351E/+*^ mice at P19 was extracted using the RNeasy Mini Kit (Qiagen, Hulsterweg, Netherland). Ribosomal RNA was depleted using a NEBNext rRNA Depletion Kit (NEB). For sequencing, a cDNA library was synthesized using the NEBNext Ultra II RNA Library Prep Kit for Illumina (NEB). Sequencing was performed on a NextSeq 500 sequencer (Illumina, San Diego, CA, USA) with 75-bp single-end reads. Resulting reads were mapped to the UCSC (University of California, Santa Cruz, CA, USA) mm10 reference genome using the Spliced Transcripts Alignment to a Reference (STAR) Aligner. The n umber of reads was calculated using RNA-Seq by Expectation Maximization (RSEM), and differentially expressed genes (DEGs) between *p62*^*+/+*^ and *p62*^*S351E/+*^ mice were analyzed by DESeq2. DEGs were visualized by volcano plots.

### Quantitative real-time PCR (qRT-PCR)

cDNAs were synthesized with 1 μg of total RNA using FastGene Scriptase Basic cDNA Synthesis (NE-LS62, NIPPON Genetics, Tokyo, Japan). qRT-PCR was performed with TaqMan® Fast Advanced Master Mix (444556, Thermo Fisher Scientific) on a QuantStudio™ 6 Pro (A43180, Thermo Fisher Scientific). Signals were normalized against *Gusb* (β-glucuronidase). Predesigned TaqMan Gene Expression Assays, including primer sets and TaqMan probes (Gusb; Mm01197698_m1, Nqo1; Mm01253561_m1, Ugdh; Mm00447643_m1 and Gstm1; Mm00833915_g1, GAPDH; Hs02786624_g1, NQO1; Hs00168547_m1, UGDH; Hs01097550_m1, GCLC; Hs00155249_m1, SOD1; Hs00533490_m1, SQSTM1; Hs00177654_m1) were purchased from Thermo Fisher Scientific.

### Statistical analysis

Statistical analyses were performed using the unpaired *t*-test (Welch *t*-test), Tukey’s test or Šidák’s multiple comparison test after one-way ANOVA. GraphPad PRISM 9 (GraphPad Software) was used for the statistical analyses. All tests were two-sided, and P values of < 0.05 were considered statistically significant.

### Resources table

| REAGENT or RESOURCE | SOURCE | IDENTIFIER |
| --- | --- | --- |
| **Antibodies** |  |  |
| Guinea pig polyclonal anti-p62/SQSTM1 (C-terminus) | Progen Biotechnik | Cat#GP62-C; RRID: AB_2687531 |
| Mouse monoclonal anti-p62 Ick ligand | BD Biosciences | Cat#610833; RRID: AB_398152 |
| Rabbit polyclonal anti-p62 (pSer403) | Gene Tex | Cat#GTX128171; RRID: AB_2885723 |
| Rabbit polyclonal anti-p62 (pSer349) | <sup>20</sup> | N/A |
| Rabbit polyclonal anti-p62 | MBL | Cat#PM045; RRID: AB_1279301 |
| Mouse monoclonal anti-p62 | Abnova | Cat#H00008878-M01; RRID: AB_1507408 |
| Rabbit polyclonal anti-KEAP1 | Proteintech | Cat#10503-2-AP; RRID: AB_2132625 |
| Rabbit monoclonal anti-ULK1 (D8H5) | Cell Signaling Technology | Cat#8054; RRID: AB_11178668 |
| Rabbit polyclonal anti-ULK2 | ABclonal | Cat#NBP-89866; RRID: AB_11009420 |
| Rabbit monoclonal anti-ATG13 (E1Y9V) | Cell Signaling Technology | Cat#13468; RRID: AB_2797419 |
| Rabbit polyclonal anti-ATG13 (pSer318) | Rockland | Cat# 600-401-C49; RRID: AB_11179920 |
| Rabbit polyclonal anti-NRF2 (H-300) | Santa Cruz Biotechnology | Cat#sc-13032; RRID: AB_2263168 |
| Rabbit polyclonal anti-RB1CC1 | Proteintech | Cat#17250-1-AP; RRID: AB_10666428 |
| Mouse monoclonal anti-β-actin | Sigma-Aldrich | Cat#A1978; RRID: AB_476692 |
| Mouse monoclonal anti-GAPDH (6C5) | Santa Cruz Biotechnology | Cat#sc-32233; RRID: AB_627679 |
| Mouse monoclonal anti-DDDDK-tag | MBL | Cat#M185-3L; RRID: AB_11123930 |
| Rabbit polyclonal anti-GFP | MBL | Cat#598; RRID: AB_591819 |
| Rabbit polyclonal anti-NQO1 | Abcam | Cat#ab9085; RRID: AB_2251526 |
| Rabbit polyclonal anti-UGDH | Abcam | Cat#ab155005 |
| Alexa Fluor Plus 488-conjugated anti-mouse IgG | Thermo Fisher Scientific | Cat#A32766; RRID: AB_2762823 |
| Alexa Fluor 488-conjugated anti-rabbit IgG | Thermo Fisher Scientific | Cat#A11034; RRID: AB_2576217 |
| Alexa Fluor Plus 594-conjugated anti-mouse IgG | Thermo Fisher Scientific | Cat#A32744; RRID: AB_2762826 |
| HRP-conjugated goat polyclonal anti-mouse IgG | Jackson ImmunoResearch Laboratories | Cat#115-035-166; RRID: AB_2338511 |
| HRP-conjugated goat polyclonal anti-rabbit IgG | Jackson ImmunoResearch Laboratories | Cat#111-035-144; RRID: AB_2307391 |
| HRP-conjugated goat polyclonal anti-guinea pig IgG | Jackson ImmunoResearch Laboratories | Cat#106-035-003; RRID: AB_2337402 |
| **Bacterial and Virus Strains** |  |  |
| *E. coli* BL21 (DE3) | Funakoshi | Cat#DS250 |
| *E. coli* DH5α | Thermo Fisher Scientific | Cat#EC0112 |
| **Chemicals** |  |  |
| Lipofectamine 3000 Transfection Reagent | Thermo Fisher Scientific | Cat#L3000015 |
| Lipofectamine RNAiMAX Transfection Reagent | Thermo Fisher Scientific | Cat#13778150 |
| Lipofectamine LTX Reagent with PLUS Reagent | Thermo Fisher Scientific | Cat#15338100 |
| DNase I | Takara Bio | Cat#2270A |
| Anti-GFP (Green Fluorescent Protein) mAb-Agarose | MBL | Cat#D153-8 |
| Glutathione Sepharose 4B | GE Healthcare | Cat#17-0756-05 |
| PreScission Protease | Cytiva | Cat#27084301 |
| One Step CBB | BIO CRAFT | Cat#CBB-1000 |
| Bafilomycin A<sub>1</sub> | KOM | Cat#BVT-0252-M001 |
| Isopropyl β-D-thiogalactopyranoside | Nacalai Tesque | Cat#19742-94 |
| cOmplete EDTA-free (protease inhibitor cocktail) | Roche Applied Science | Cat#05056489001 |
| PhosSTOP (phosphatase inhibitor cocktail) | Roche Applied Science | Cat#4906845001 |
| Polyvinylpyrrolidone | Sigma-Aldrich | Cat#PVP10 |
| **Experimental Models: Cell Lines** |  |  |
| Huh-1 | RIKEN | Cat#JCRB0199; RRID: CVCL_2956 |
| Huh-1 *p62* KO | <sup>20</sup> | N/A |
| Huh-1 *FIP200* KO | This study | N/A |
| Huh-1 *p62*, *KEAP1* DKO | This study | N/A |
| Huh-1 *ULK1* KO | This study | N/A |
| HEK293T | ATCC | Cat#CRL-3216 |
| PlatE | Cell BIOLABS | Cat#RV-101 |
| **Experimental Models: Organisms/Strains** |  |  |
| Mouse: *p62*<sup>S351A/+</sup> | This study | N/A |
| Mouse: *p62*<sup>S351A/S351A</sup> | This study | N/A |
| Mouse: *p62*<sup>S351E/+</sup> | This study | N/A |
| **Recombinant DNA** |  |  |
| pGEX6P-1 | GE Healthcare | #28954648 |
| pGEX6P-1-8xUb | Genscript | N/A |
| pGEX6P-KEAP1 | This study | N/A |
| pGEX6P-mCherry-p62 | This study | N/A |
| pGEX6P-mCherry-p62<sup>S349E</sup> | This study | N/A |
| pGEX6P-mCherry-p62<sup>S403E</sup> | This study | N/A |
| pEGFP-C2 | Clontech | Cat#6083-1 |
| pEGFP-C2-p62 | This study | N/A |
| pEGFP-ULK1 | This study | N/A |
| pRetroX-TetOne-Puro-EGFP-p62 | This study | N/A |
| pRetroX-TetOne-Puro-EGFP-p62<sup>S349E</sup> | This study | N/A |
| pRetroX-TetOne-Puro-EGFP-p62<sup>S349A</sup> | This study | N/A |
| pRetroX-TetOne-Puro-EGFP-p62<sup>T350A</sup> | This study | N/A |
| pIRES-puro-N3xFLAG | This study | N/A |
| pIRES-puro-N3xFLAG-ULK1 | This study | N/A |
| pIRES-puro-N3xFLAG-ULK2 | This study | N/A |
| **Software and Algorithms** |  |  |
| ImageJ | Thermo Fisher Scientific | RRID: SCR_014477 |
| CellPath Finder | <sup>56</sup> | RRID: SCR_002285 |
| Evolution-Capt Edge | Yokogawa | N/A |
| FV10-ASW 04.01 | Vilber Bio Imaging | N/A |
| Multi Gauge (version: 3.4) | Fuji Film | N/A |
| FV31S-SW (version: 2.4.1.198) | Olympus | N/A |
| Kodec | Olympus | N/A |
| Biomolecular AFM viewer-2.1 | <sup>54</sup> | N/A |
| GraphPad Prism 9 | <sup>57</sup> | N/A |
| Adobe Photoshop CS6 | Adobe | N/A |

## Acknowledgements

We thank Dr. Mikako Shirouzu for providing purified p62 (268-440aa) and p62 (320-440aa) proteins and Ms. Miyako Yoshimura for technical assistance. We also thank Mr. Katsuyuki Kanno, and Mr. Takayuki Yabe for their help with CLEM and histological analyses. D.N. is supported by a Grant-in-Aid for Young Scientists (19K16344) and the Takeda Science Foundation. H.M. is supported by a Grant-in-Aid for Scientific Research (B) (20H03213), by Advanced Research and Development Programs for Medical Innovation (AMED-PRIME, 21gm6410019h0001), and by the Inamori foundation. S.K., T.F., and R.A. are supported by Grants-in-Aid for Scientific Research (C) (20K06549 for S.K., 21K06178 for T.F., and 22K06300 for R.A.). S.K-H. is supported by a Grant-in-Aid for Encouragement of Scientists (21H04163). S.W. is supported by a Grant-in-Aid for Scientific Research (B) (20H03415). Y.I. is supported by a Grant-in-Aid for Scientific Research (C) (20K06644). N.N.N. is supported by a Grant-in-Aid for Scientific Research on Innovative Areas (19H05707), JST CREST (JPMJCR20E3) and by the Platform Project for Supporting Drug Discovery and Life Science Research (Basis for Supporting Innovative Drug Discovery and Life Science Research (BINDS)) from AMED under grant no. JP19am0101001 (support no. 0002). M.K. is supported by a Grant-in-Aid for Scientific Research on Innovative Areas (19H05706), a Grant-in-Aid for Scientific Research (A) (21H004771), the Advanced Research and Development Programs for Medical Innovation (AMED-CREST, 22gm1410004h0003), the Japan Society for the Promotion of Science (an A3 foresight program), and the Takeda Science Foundation.

## Author Contributions

Y.I., N.N.N., and M.K. designed and directed the study. R.I., S.K., H.M., S.T., T.F., S.K-H., and Y.I. carried out the biochemical and cell biological experiments. S.K-H. generated knockout cell lines. D.N, Y.F., and N.N.N conducted high-speed AFM and in vitro LLPS analysis. R.A., E.L., and S.W. carried out histological analysis of mice. T.K. and M.N. performed and analyzed RNA-seq analysis. H.A. provided intellectual support. N.N.N. and M.K. wrote the manuscript. All authors discussed the results and commented on the manuscript.

## Figure legends

**Supplementary Fig. S1 HS-AFM observation of SNAP-ULK1 and p62 (268–440 aa), and complex of SNAP-Atg1/p62 (268–440 aa)**

**a**,**b** Successive HS-AFM images of SNAP-ULK1 (a) and p62_268–440 (b). Height scale: 0– 4.4 nm (a), 0–3.4 nm (b); scale bar: 20 nm (a,b). **c** Successive HS-AFM images of p62_268– 440 with SNAP-Atg1. Height scale: 0–3.6 nm; scale bar: 30 nm. **d** Schematics showing the molecular characteristics determined by HS-AFM. Grey spheres, globular domains consisting of N-terminal KD and C-terminal MIT of Atg1; pink spheres, globular domains consisting of C-terminal UBA domain of p62; blue thick solid lines, IDRs.

**Supplementary Fig. S2 Generation of *ULK1* knockout**, ***FIP200* knockout, and *p62*/*KEAP1* double-knockout cell lines**

Immunoblot analysis. The indicated genotype cell lines were lysed, then subjected to SDS-PAGE followed by immunoblot analysis with the indicated antibodies. The asterisk indicates non-specific bands.

**Supplementary Fig. S3 An mTORC1 inhibitor has no effect on the phosphorylation level of p62 in Huh1**

Immunoblot analysis. Huh-1 cells were treated with 250 nM Torin-1 for 6 h, and the cell lysates were subjected to immunoblot analysis with the indicated antibodies. The asterisk indicates non-specific bands. Data shown are representative of three or four separate experiments. Bar graphs show the results of quantitative densitometric analysis of Ser349-or Ser403-phosphorylated p62 forms relative to total p62 (*n* = 4), Thr389-phosphorylated p70S6K relative to total p70S6K (*n* = 3), and Ser318-phosphorylated ATG13 relative to total ATG13 (*n* = 4). Data are means ± s.e. Statistical analysis was performed by Šidák’s test after one-way ANOVA.

**Supplementary Fig. S4 Effect of ULK1 and the ULK2 inhibitor ULK-101 on p62 phosphorylation**

Immunoblot analysis. Huh-1 cells were treated with or without 7.5 μM ULK-101 for 6 h, and the cell lysates were subjected to immunoblot analysis with the indicated antibodies. The asterisk indicates non-specific bands. Data shown are representative of three separate experiments. Bar graphs show the results of quantitative densitometric analysis of S349- or S403-phosphorylated p62 forms relative to total p62 (*n* = 3), and of S318-phosphorylated ATG13 relative to total ATG13 (*n* = 3). Data are means ± s.e. Statistical analysis was performed by Welch’s *t*-test.

**Supplementary Fig. S5 *In vitro* formation of p62**^**S349E_S403E_S407E**^**-8xUb condensates**

10 μM mCherry-p62^S349E_S403E_S407E^ was mixed with 10 μM SNAP-8xUb labeled with SNAP-Surface 649. Scale bar: 20 μm.

**Supplementary Fig. S6 Gross anatomy of the stomach of *p62***^***+/+***^ **and *p62***^***S351E/+***^ **mice** The forestomach of *p62*^*S351E*^ heterozygotes was obviously thickened compared with that of wild-type mice.

**Supplementary Fig. S7 Analysis of mice lacking p62 phosphorylation at S351**

**a** External appearance of *p62*^*+/+*^, *p62*^*S351A/+*^, and *p62*^*S351A/S351A*^ mice at postnatal day (P) 19. **b, c** Body weight (g) (**b**) and liver weight (% of body weight) (**c**) of *p62*^*+/+*^ (*n* = 3), *p62*^*S351A/+*^ (*n* = 6), and *p62*^*S351A/S351A*^ mice (*n* = 3) at P19. Data are means ± s.e. Statistical analysis was performed by Tukey’s test after one-way ANOVA. **d** Gene expression of NRF2 targets. Total RNAs were prepared from mouse livers of *p62*^*+/+*^ (*n* = 3), *p62*^*S351A/+*^ (*n* = 6), and *p62*^*S351A/S351A*^ mice (*n* = 3) at P19. Data are means ± s.e. Statistical analysis was performed by Tukey’s test after one-way ANOVA. **e** Immunoblot analysis of *p62*^*+/+*^ (*n* = 3), *p62*^*S351A/+*^ (*n* = 6), and *p62*^*S351A/S351A*^ mice (*n* = 3) at P19. Liver homogenates were subjected to immunoblot analysis with the indicated antibodies. Bar graphs show the results of quantitative densitometric analysis. Data are means ± s.e. Statistical analysis was performed by Tukey’s test after one-way ANOVA. **f** Hematoxylin and eosin (HE) staining of livers from *p62*^*+/+*^, *p62*^*S351A/+*^, and *p62*^*S351A/S351A*^ mice at P19. Scale bar, XX μm. **g** Serum levels of aspartate aminotransferase (AST), alanine aminotransferase (ALT), glucose, total cholesterol, blood urea nitrogen (BUN), and creatinine from *p62*^*+/+*^ (*n* = 3), *p62*^*S351A/+*^ (*n* = 6), and *p62*^*S351A/S351A*^ mice (*n* = 3) at P19 were measured. IU/l, international units/liter. Data are means ± s.e. Statistical analysis was performed by Tukey’s test after one-way ANOVA.

**Supplementary Movie S1**

HS-AFM movie of SNAP-ULK1. The images were acquired at 5 fps. Height scale: 0–3 nm. Scale bar: 20 nm.

**Supplementary Movie S2**

HS-AFM movie of p62_268–440. The images were acquired at 5 fps. Height scale: 0–3 nm. Scale bar: 20 nm.

**Supplementary Movie S3**

HS-AFM movie of p62_268–440 with SNAP-ULK1. The images were acquired at 5 fps. Height scale: 0–3 nm. Scale bar: 20 nm.

**Supplementary Movie S4**

HS-AFM movie of p62_268–440 with SNAP-Atg1. The images were acquired at 5 fps. Height scale: 0–3 nm. Scale bar: 30 nm.

**Supplementary Movie S5**

Representative time-lapse image of mCherry-KEAP1 after photobleaching of whole GFP-p62^S349E^ body (Scale bar: 2 μm).

**Supplementary Movie S6**

Representative time-lapse image of mCherry-KEAP1 after photobleaching of whole GFP-p62^S349A^ body (Scale bar: 2 μm).

**Supplementary Movie S7**

Representative time-lapse image showing fluorescence loss of mCherry-KEAP1 in GFP-p62^S349E^ bodies after photobleaching over a large area of cells (Scale bar: 20 μm).

**Supplementary Movie S8**

Representative time-lapse image of mCherry-KEAP1. The fluorescence loss of mCherry-KEAP1 in GFP-p62^S349A^ bodies after photobleaching at a large area of cells (Scale bar: 20 μm).

**Supplementary Movie S9**

Representative time-lapse image of mCherry-KEAP1 after photobleaching of the central portion of p62^S349E^ body (Scale bar: 2 μm).

**Supplementary Movie S10**

Representative time-lapse image of mCherry-KEAP1 after photobleaching of the central portion of p62^S349A^ body (Scale bar: 2 μm).

